# Inferring tumor-specific cancer dependencies through integrating ex-vivo drug response assays and drug-protein profiling

**DOI:** 10.1101/2022.01.11.475864

**Authors:** Alina Batzilla, Junyan Lu, Jarno Kivioja, Kerstin Putzker, Joe Lewis, Thorsten Zenz, Wolfgang Huber

**Affiliations:** Genome Biology Unit, European Molecular Biology Laboratory (EMBL), Heidelberg, Germany; Medical Faculty, Heidelberg University, Heidelberg, Germany; Faculty of Biosciences, Heidelberg University, Heidelberg, Germany; Molecular Medicine Partnership Unit (MMPU), Heidelberg, Germany; Department of Medical Oncology and Hematology, University Hospital Zürich and University of Zürich, Zürich, Switzerland; Chemical Biology Core Facility, European Molecular Biology Laboratory (EMBL), Heidelberg, Germany

## Abstract

The development of cancer therapies may be improved by the discovery of tumor-specific molecular dependencies. The requisite tools include genetic and chemical perturbations, each with its strengths and limitations. Drug perturbations can be readily applied to primary cancer samples at a large scale, but mechanistic understanding of hits and further pharmaceutical development is often complicated by the fact that a small compound has a range of affinities to multiple proteins.

To computationally infer the molecular dependencies of individual cancers from their ex-vivo drug sensitivity profiles, we developed a mathematical model that deconvolutes these data using measurements of protein-drug affinity profiles.

Our method, DepInfeR, correctly identified known dependencies, including EGFR dependence in Her2+ breast cancer cell line, FLT3 dependence in AML tumors with FLT3-ITD mutations and the differential dependencies on the B-cell receptor pathway in two major subtypes of chronic lymphocytic leukemia (CLL). Furthermore, our method uncovered new subgroup-specific dependencies, including a previously unreported dependence of high-risk CLL on Checkpoint kinase 1 (CHEK1). The method also produced a more accurate map of the molecular dependencies in a heterogeneous set of 117 CLL samples.

The ability to deconvolute polypharmacological phenotypes into underlying causal molecular dependencies should increase the utility of high-throughput drug response assays for functional precision oncology.

## Introduction

The sustained proliferation or apoptosis resistance of cancer cells rely on abnormal pathway activities, often resulting from genetic or epigenetic alterations. The molecular processes underlying these activities, however, can vary greatly across cancer types, and even between tumors from different patients with the same cancer type. Such heterogeneity of driver dependencies can lead to differential treatment responses. A key aim of precision cancer therapy is to exploit core vulnerabilities of each individual tumor.

A powerful, scalable, and systematic approach to identify cancer-specific dependencies uses genetic perturbations (McFarland *et al*, 2018; Behan *et al*, 2019; Cowley *et al*, 2014; Tsherniak *et al*, 2017) such as RNAi and CRISPR/Cas9 systems to knock down or knock out a specific gene in cancer cells and observe the subsequent effect on cell growth or survival. While RNAi and CRISPR/Cas9 screens can be applied for nearly any gene in cell line models, their use remains challenging in primary tumor samples (Laustsen & Bak, 2019; Mandal *et al*, 2014), which poses an important limitation for clinical research. This is the case especially for indolent diseases, such as chronic lymphocytic leukemia (CLL). Furthermore, significant discrepancies may exist between the effects of a genetic perturbation and targeted drug inhibition of the encoded protein (Gonçalves *et al*, 2020). This is potentially because a drug might quantitatively inhibit the enzymatic activity of a protein but leave other functions, such as scaffolding, unaffected (Weiss *et al*, 2007), whereas the genetic perturbation completely disrupts all functions. Off-target effects (Morgens *et al*, 2017) and side effects, such as inducing cellular DNA damage response (Fellmann *et al*, 2017; Munoz *et al*, 2016), of the CRISPR/Cas9 system may further complicate analysis.

A major goal of cancer dependency mapping is to propose druggable targets for cancer treatment. Chemical perturbation experiments, which rely on high-throughput screening of bioactive compounds on cancer cells, provide an attractive complementary approach to genetic perturbations due to their high translational value for personalized oncology (Tyner *et al*, 2018; Yang *et al*, 2012). Chemical perturbations are readily applicable to primary tumor models, and thus can identify patient- and tumor-specific dependencies. Drug treatment of primary tumor cells also better approximate the ex-vivo cellular effects. By design, all identified protein dependencies are “targetable”. However, a major challenge is the polypharmacological nature of small molecular compounds, as many of them show a broad range of off-target effects, which obstructs the identification of the protein targets related to the desired phenotype (Klaeger *et al*, 2017).

To improve the utility of high-throughput drug sensitivity data for functional precision medicine, we developed a computational method that integrates two experimentally accessible input data matrices: the drug sensitivity profiles of cancer cell lines or primary tumors ex-vivo (X), and the drug affinities of a set of proteins (Y), to infer a matrix of molecular protein dependencies of the cancers 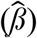. We deconvolve the protein inhibition effect on the viability phenotype by using regularized multivariate linear regression. This framework assigns an “dependence coefficient” to each protein and each sample, and therefore could be used to gain causal and accurate understanding of functional consequences of genomic aberrations in a heterogenous disease, as well as to guide the choice of pharmacological intervention for a specific cancer type, sub-type, or a patient. We termed our computational framework as DepInfeR and implemented it as an R package (https://github.com/Huber-group-EMBL/DepInfeR).

As similar ex-vivo assays can be readily performed on cell lines and primary samples, with established platforms and ready to use datasets for both, drug sensitivity profiling (Lu *et al*, 2021; Dietrich *et al*, 2017; Iorio *et al*, 2016; Yang *et al*, 2012) and drug-protein profiling (Klaeger *et al*, 2017), we expect our approach to be adaptable to different diseases.

## Material and Methods

### Processing of the drug response assay datasets

#### The Genomics of Drug Sensitivity in Cancer (GDSC) Project data

the GDSC1 dataset of drug sensitivities was downloaded from the Genomics of Drug Sensitivity in Cancer Project (https://www.cancerrxgene.org; (Yang *et al*, 2012), separately for different cancers: diffuse large B-cell lymphoma (DLBCL, n=30), acute lymphocytic leukemia (ALL, n=25), acute myeloid leukemia (AML, n=24) and breast carcinoma (BRCA, n=47). As a drug sensitivity measure, the dataset includes the area under the curve (AUC), defined as the area under the sigmoid-fit dose-response curve (5-point or 9-point) (Vis *et al*, 2016). The AUC values were converted into *z*-scores, by computing mean and standard deviation for each cell line, across the drugs. These *z*-scores were used as the drug sensitivity matrix *Y*, including sensitivity data for 66 drugs. Cell lines with more than 24 (≙ 35%) missing values were removed, resulting in a total of 126 cell lines. The missForest imputation method from the missForest package (*V 1*.*4*; Stekhoven & Bühlmann, 2011) was used to impute missing values in the drug sensitivity matrix for the remaining cell lines.

The cancer cell lines were annotated with their cancer types and mutation status obtained from the GDSC project page. Additionally, Her2 status of breast cancer cell lines were annotated based on information from prior publications (Dai *et al*, 2017; Jernström *et al*, 2017).

#### BeatAML data

the BeatAML dataset (Tyner *et al*, 2018) contains ex-vivo drug sensitivity data of 528 tumor specimens collected from acute myeloid leukemia (AML) patients. The data were downloaded from the National Cancer Institute’s CTD2 Network (BeatAML_Waves1_2). The AUC from a 7-point dose-response curve was used as described by Tyner *et al*. 2018 and the *z*-scores of the AUC for each cell line across all drugs were used as the response value for generating the drug sensitivity matrix *Y*, including sensitivity data for 61 drugs. Samples with more than 15 (≙ 24%) missing values were removed, resulting in a total number of 421 samples. Cut-off selection and missing value imputation were done as described for the GDSC1 dataset.

#### Ex-vivo drug response assay on primary leukemic tumor samples

we performed ex-vivo drug response profiling on primary tumor samples from CLL (n=117), mantle cell lymphoma (MCL, n=7), or T-cell prolymphocytic leukemia (T-PLL, n=7) patients. We termed this drug screen dataset as the EMBL2016 dataset. The samples were annotated with available genomic patient metadata. To obtain a single drug by tumor samples matrix, relative cell viabilities (compared to treatment with DMSO control) were averaged from 9 drug concentrations. Subsequently, the z-score of these values for each patient across all drugs was used as the response value for generating drug sensitivity matrix *Y*, including sensitivity data for 85 drugs.

### Processing of the kinase inhibitor profiling data

The kinase inhibitor profiling data were obtained from Supplementary Table S2 of *Klaeger et al. 2017*. The data were subset for drugs overlapping with those used in either of the drug response assays described above by matching the drug names. The drug synonym annotation (Klaeger *et al*, 2017) was considered and naming variations were manually curated. The data were arranged in a matrix *X* of dissociation constants (*K*_*d*_) for each drug-protein pair. The *K*_*d*_ values were transformed with a sigmoid function into the range [0, 1]. We used *f*(*x*) = (arctan((-log_10_(*x*) + 2) * 3) + *π*/2) / *π*, with x being the raw *K*_*d*_ values. Values of *f(x)* close to 1 correspond to evidence of drug-protein binding, values close to 0 to absence of such evidence. The graph of the function *f* and the distribution of values before and after the transformation are shown in Supplementary Figure S1.

As some kinases had similar ligand profiles, and since this drug-target matrix was intended to be the design matrix in a multivariate regression model, highly similar kinases were “collapsed” in order to achieve model identifiability and parameter estimate stability. To this end, we computed the cosine similarity between each pair of kinases, applied hierarchical clustering, and identified subtrees in the cluster dendrogram in which all kinases had a cosine similarity > 0.8. For each such cluster, the kinase with the largest sum of transformed *K*_*d*_ values within the cluster was used as a cluster name, while the other kinases were also recorded and retained for case-by-case follow-up. The collapsed kinase sets are provided in Supplementary Figure S2. For the analysis of the patient primary cell assays, the drug-protein profiling matrix was additionally filtered by removing kinases that were not, or only weakly expressed in the tumor samples, as follows. In the gene-level count matrix of the RNA sequencing data of the tumor samples, the 80% quantile of its counts across samples was computed for each gene, and a gene was considered not or only weakly expressed if this value was less than 10. Thus, 20 of the 377 kinases were removed from the downstream analysis for the EMBL2016 dataset, and 14 of the 380 kinases in the BeatAML analysis. The final dimensions of the input matrices X were: GDSC: 66 drugs × 118 proteins; BeatAML: 61 drugs × 112 proteins; EMBL2016: 85 drugs × 131 proteins (Figure 1A).

**Figure 1.**
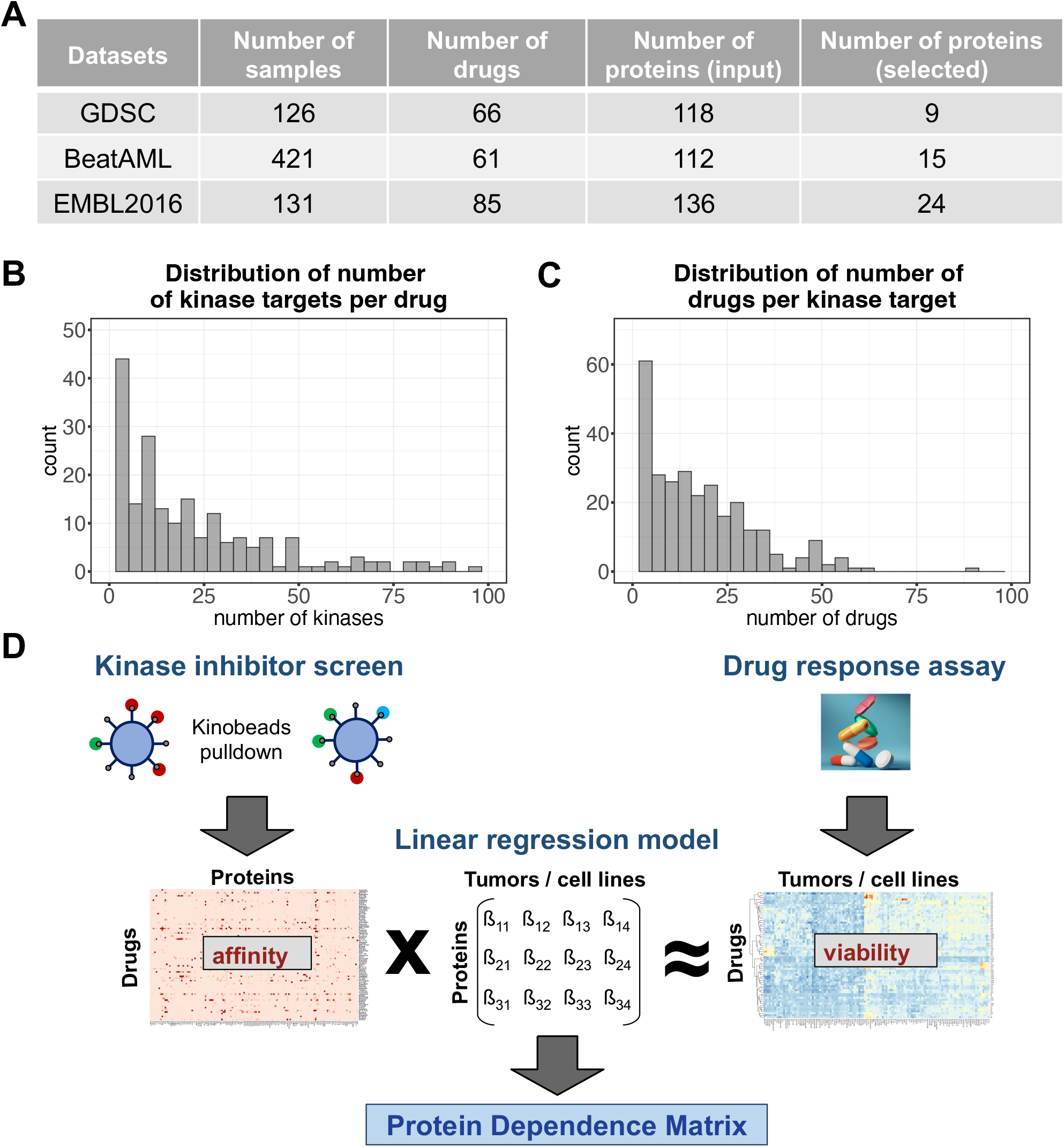
Principle of the protein dependence inference framework. (**A**) Summary of tumor samples and drugs in the three datasets used in this study, and the number of proteins with non-zero coefficients in the inferred protein dependence matrix. (**B**) Distribution of the number of kinases bound by each drug (*k*_*d*_ < 1000nM) and (**C**) distribution of the number of drugs binding each kinase from the kinobeads kinase profiling data used in our analysis. (**D**) Inference of the protein dependence matrix using a multivariate multi-response regression model with L1 regularization. A known drug-protein affinity matrix (independent variables) and a known drug-effect matrix (response variables) are used as input for the model to infer the unknown protein dependence matrix.

### Regularized multivariate regression model

Regularized multivariate regression was used to obtain the protein dependence coefficients. We fit the model

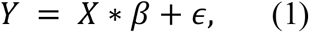

where *X* is the processed drug-protein binding matrix (drugs x proteins), Y is the processed drug sensitivity matrix (drugs x samples), β is the unknown protein dependence matrix (samples x proteins), and ε is a matrix of residuals (drugs x samples). Given the data for *X* and *Y*, we computed the model fit for β by minimising ε using a multi-response Gaussian linear model (family = “mgaussian”) with L1-penalty (lasso) on β using the *glmnet* package (*V 3*.*0-2;* (Friedman *et al*, 2010)) with the mixing parameter *α*=1, and λ shared among samples.

We repeated the following 100 times and computed the element-wise medians of the 100 β-matrices to obtain the final estimate 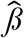: determine an optimal penalty parameter λ that minimizes the cross-validated mean squared error using the function *cv*.*glmnet* from the glmnet package with 3-fold cross-validation and fit the model with the optimal choice of λ. As a quality measure, the coefficient of determination R^2^ from the final ß estimate was used.

### Principal Component Analysis (PCA) visualization

PCA analysis of 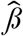 was performed using the R function *prcomp* from the *stats* package, which centers and scales the columns (corresponding to samples) of the matrix for equal variance. Visualization graphics were made using the *factoextra* package (*V 1*.*0*.*6*; (Kassambara & Mundt, 2019)).

### *k*-means clustering and Rand index calculation

*k*-means clustering analysis was performed on 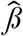 using the *eclust* function from the *factoextra* package, which centers and scales the columns of the matrix. Class information (five cancer types in the case of the GDSC data, IGHV mutational status for the EMBL2016 CLL data) was withheld from the clustering method, but used to set the number of clusters *k*. 50 different random starting configurations (*nstart*) were attempted within the *eclust* function and the best configuration was chosen. As a measure of concordance of the resulting clusters and the ‘ground truth’ classes, the adjusted Rand Index (Rand, 1971) was calculated using the *cluster*.*stats* function from the *fpc* package (*V 2*.*2-9;* (Hennig & Imports, 2015)). *k*-means clustering of the drug sensitivity matrix was performed analogously.

### Differential gene expression analysis

To identify gene expression signatures related to IGHV mutation status, an RNA sequencing dataset from the European Genome-phenome Archive (EGA; http://www.ebi.ac.uk/ega/, accession: EGAS00001000374 (Ferreira *et al*, 2014)) was analyzed. Differential expression testing was performed using the *DESeq2* package (*V 1*.*28*.*1*; (Love *et al*, 2014)). Genes with overall low counts (sum across samples < 10) were excluded from the analysis. The *camera* method was used to test for differential expression of hallmark gene sets from the MSigDB database v7.2 (Subramanian *et al*, 2005; Liberzon *et al*, 2015), using the implementation in the eponymous function in the *limma* package (*V 3*.*44*.*3*; (Ritchie *et al*, 2015).

To identify gene expression signatures related to B-cell stimulation, we analyzed gene expression microarray dataset of anti-IgM triggering in primary CLL samples (E-GEOD-39411) (Vallat *et al*, 2013). The *limma* package was used to perform variance stabilizing normalization and differential expression testing, and the *camera* method was used to test for differential expression of Hallmark gene sets as above.

### Other statistical methods

Association tests were performed using Student’s *t*-test (two-sided, with equal variance) and two-way ANOVA. The Benjamini-Hochberg method was applied to the *P* values to account for multiple testing.

### Data and code availability

All data and code used in this study are available at (https://github.com/Huber-group-EMBL/DepInfeR_workflow). A website of rendered html reports is available at: https://www.huber.embl.de/users/jlu/depInfeR/index.html We provide an R/Bioconductor package, DepInfeR (https://github.com/Huber-group-EMBL/DepInfeR), for users to estimate cancer protein dependence coefficient using their own datasets.

## Results

We aimed to infer the extent and direction of the dependence of individual tumor samples on individual proteins. To perform this, we used two available data types (Fig. 1A): 1) a drug-protein profiling matrix, which reports binding affinities of each of a set of drugs to proteins, and 2) a drug sensitivity matrix, which reports the effects on cell viability of each drug across a set of tumor samples. The effects are measured as dimensionless ratios, relative cell viability compared to DMSO (dimethyl sulfoxide) control.

As the first data source, we used the kinase inhibitor profiling dataset by *Klaeger et al*., *2017*. The authors immobilized 243 kinase inhibitors on kinobeads and detected presence and abundance of kinases binding after exposition of the beads to cell lysate by mass spectrometry proteomics. They reported their results in terms of *K*_*d*_ values, which we converted into binding scores in the range between 0 and 1 (Methods). The data show a large variety in the degree of selectivity among kinase inhibitors: while some bind selectively to only a handful of kinases, many bind dozens of different cellular proteins (Fig. 1B). Similarly, many kinases are bound by multiple drugs (Fig. 1C). Altogether, the picture implied by the data of *Klaeger et al*., *2017* represents an intricate, spread out network of drug-kinase binding with few one-to-one pairs or clear drug-kinase clusters (Supplementary Figure S3).

The objective of our approach is to use this network to infer the kinases that a particular sample depends on for viability by overlaying the network with the effects of the drugs on the sample. This task can be posed in terms of multivariate multi-response regression involving three matrices: the known drug-protein binding matrix, a known drug-sample sensitivity matrix -below we consider three different instances of such a matrix - and the unknown protein dependence matrix (Fig. 1A). While no useful inference can be expected if there is only data for a single or a small number of samples, the regression becomes increasingly identifiable as the number of samples increases. The matrix of fitted coefficient values indicates to what extent the viability of each sample depends on each protein. High positive coefficients imply strong reliance of a particular sample on a particular protein for survival, and therefore a high vulnerability towards inhibition of the protein. Conversely, proteins with coefficients close to zero are less essential for the cell’s immediate survival. Negative coefficients indicate that the viability phenotype benefits from inhibition of the protein. As the regression problem is high-dimensional, we used L1 (LASSO) regularization with cross-validation to achieve parsimonious and stable regression results and select the most predictive features. Moreover, we collapsed sets of nearly collinear independent variables (i.e., proteins with highly similar drug binding profiles) into groups represented by one of the variables (Supplementary Figure S2). We termed our method as DepInfeR and implemented an R package with the same name to perform the data preprocessing, the multivariate regression and various downstream analyses and visualizations.

As the second data source, we used three separate drug sensitivity datasets: one from cancer cell line collections and two from drug sensitivity studies of primary blood cancer samples (AML and CLL).

### Inferred protein dependence profiles reflect differential inter- and intra-cancer type dependencies in cell lines

As a first test of DepInfeR, we applied it to the cancer cell line drug sensitivity data from the GDSC database (Yang *et al*, 2012). We used the subset of data for cell lines derived from diffuse large B-cell lymphoma (DLBCL), acute lymphocytic leukemia (ALL), acute myeloid leukemia (AML), and breast carcinoma (BRCA). We concatenated these data into a single matrix of relative cell line viabilities (n=126) under the effect of the different drugs (n=66). Besides the relative viabilities, the matrix did not contain explicit information on cancer type or any other properties of the cell lines. We inferred the protein dependence matrix and visualized it with a heatmap with rows and columns ordered in accordance with hierarchical clustering (Supplementary Figure S4). The four different cancer types were separated in these visualizations, indicating that they have distinctive protein dependence profiles (Fig. 2A, B). The breast cancer cell lines further split into two groups, which corresponded to their Her2 status—an important prognostic marker based on the assessment of the expression level of the epidermal growth factor receptor 2 (Her2 / EGFR) (Slamon *et al*, 1987; Ahn *et al*, 2020). DepInfeR correctly detected the dependence of Her2-positive breast cancer cell lines on EGFR (Epidermal Growth Factor Receptor) signaling (Fig. 2A, B). Irrespective of their Her2 status, the breast cancer cell lines showed higher vulnerability to the inhibition of JAK1 (Janus Kinase 1) than the leukemia cell lines (Fig. 2A). ALL and AML cell lines showed a significantly higher dependence on the receptor tyrosine kinase FLT3 (Fig. 2A), in line with the known importance of mutated FLT3 in AML and ALL (Annesley & Brown, 2014; Griffith *et al*, 2016; Armstrong *et al*, 2004).

**Figure 2.**
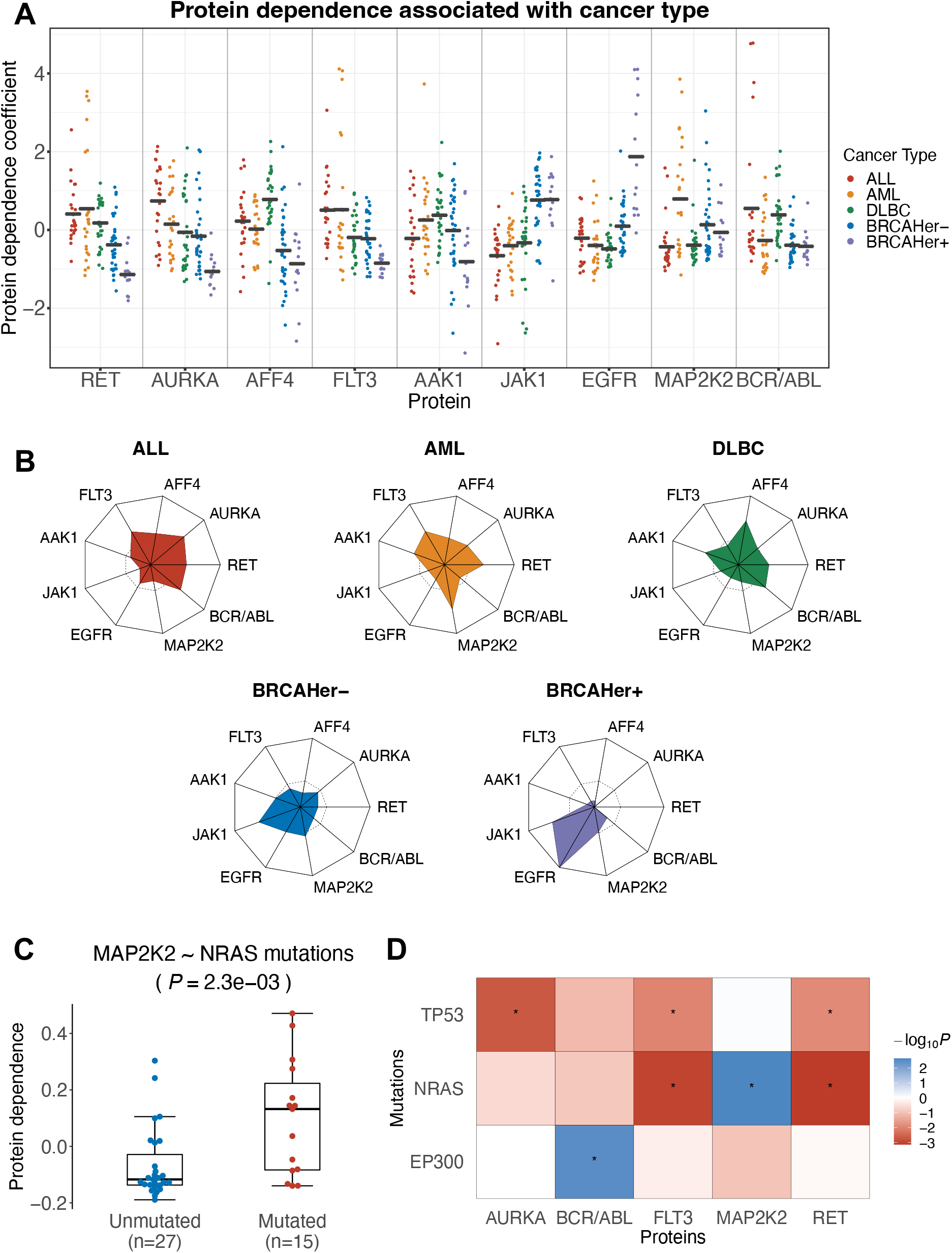
Results of protein dependence inference on the GDSC1 dataset. (**A**) Shown are the data for 9 kinases for which we found significant differences in the protein dependence coefficients between cancer types (ANOVA, BH adjusted p-value < 0.05, Fold Change > 0.1). Within each panel, each point corresponds to a cell line. The coefficients were centered and scaled to obtain a per-protein z-score, and the points are grouped and colored by cancer type (ALL, red; AML, orange; DLBC, green, BRCAHer-, blue; BRCAHer+, purple). (**B**) Radar plots of protein dependence coefficients across the different cancer types. Dashed line represents a protein dependence coefficient of 0. (**C**) Association between *NRAS* mutation status and MAP2K2 dependence. Association testing was performed using Student’s t-test (two-sided, with equal variance) or ANOVA test and Benjamini-Hochberg correction was applied. (**D**) A heatmap showing the -log10(*P*-values) with signs determined by direction of fold changes of the associations between mutational background of the cell lines and protein dependence coefficients (Student’s t-test). Blue: associations with higher dependence coefficients in mutated cases; red: lower dependence coefficients in mutated cases; stars indicate the associated pass 10% FDR control.

DepInfeR identified further, more subtle protein dependence profile differences that were associated with genetic aberrations of cell lines within one cancer type (Fig. 2C). *NRAS* mutation status was associated with the dependence coefficient of multiple kinases. Its positive association with MAP2K2 (Fig. 2D) reflects the higher activity of the MAPK/ERK kinase pathway in tumors driven by *NRAS* mutation (Raaijmakers *et al*, 2016). Conversely, decreased sensitivity of *NRAS* mutated cell lines to the inhibition of the tyrosine-protein kinases RET and FLT3 (Fig. 2C) is consistent with the fact that these two proteins are upstream activators of the RAS pathway: their inhibition is expected to be of little consequence if RAS is constitutively activated by *NRAS* mutation (McMahon *et al*, 2019; Jhiang, 2000; Köthe *et al*, 2013).

### Inferring cancer dependencies using ex-vivo drug sensitivity profiling on primary leukemic samples

#### DepInfeR method correctly infers kinase dependencies in AML

We next applied DepInfeR to a dataset of primary tumor samples, the BeatAML data, which report the ex-vivo drug sensitivities of 672 tumor specimens from 562 AML patients (Tyner *et al*, 2018). The inferred protein dependence coefficients indicated significantly higher dependence on FLT3 for tumors with *FLT3*-ITD mutation, a known proliferative driver in AML (Daver *et al*, 2019), consistent with the findings by Tyner et al. and in line with biological expectation (Supplementary Figure S5). An additional positive correlate of *FLT3*-ITD was the inferred dependence on the Lck kinase. This association was not described by *Tyner et al. 2018*, but supports cooperative roles of Lck activity with *FLT3*-ITD mutation as suggested previously (Marhäll *et al*, 2017). We also identified the vulnerability of *KRAS* and *NRAS* mutated tumors to MAPK-pathway inhibition (Supplementary Figure S5).

#### The protein dependence profiles of primary CLL samples recapitulate major molecular subgroups

Having established the feasibility of protein dependence inference on these existing datasets, we proceeded to generate a dataset tailored to the study of lymphocytic leukemias. We selected samples from 131 patients diagnosed with three prevalent leukemia types: CLL (n=117), MCL (n=7), and T-PLL (n=7). This study design enabled us to consider entity specific properties of MCL and T-PLL, and to further resolve the considerable intra-entity heterogeneity of CLL with respect to molecular factors including the mutational status of immunoglobulin heavy chain variable region (IGHV) genes, trisomy 12 status, and *TP53* mutations (Dietrich *et al*, 2017; Zenz *et al*, 2010). We measured drug sensitivity phenotypes of these tumors towards 85 kinase inhibitors and applied DepInfeR to infer the protein dependence map (Supplementary Figure S6).

We observed that kinases from the cell cycle regulation machinery, such as CDK6, CDK17, and CCNT1, had uniformly high dependence coefficients across samples, reflecting the high toxicity of inhibitors targeting cell cycle (Supplementary Figure S7). By contrast, kinases from specific signaling pathways, such as the Bruton’s tyrosine kinase (BTK) from the B cell receptor (BCR) pathway, showed varied coefficients among samples (Supplementary Figure S7).

The T-PLL samples were clearly separated from CLL and MCL by their characteristically low dependence on the BTK (Supplementary Figure S7), which is consistent with the role of BTK as a main pathway component of BCR signaling and major driver of B-cell leukemias (Hendriks *et al*, 2014).

Next, we zoomed in to study the relationship between the molecular heterogeneity of our CLL samples with the protein dependencies. In a hierarchical clustering of the samples based on their protein dependence profiles, the most striking pattern was the separation between CLLs with mutated IGHV status (M-CLL) and unmutated IGHV status (U-CLL) (Supplementary Figure S6). The IGHV mutational status reflects the cell of origin in CLL and remains an important prognostic factor: U-CLL shows significantly faster disease progression and worse clinical outcome than M-CLL (Damle *et al*, 1999).

Clustering and other forms of pattern detection on the samples can, in addition to the protein dependence map, also be done on the drug sensitivity data directly. We aimed to compare the two approaches by assessing the results of k-means clustering with the Rand index (Rand 1971). We chose k=2 in order to differentiate between U-CLL and M-CLL. While both approaches broadly separated the two subgroups, clustering based on the protein dependence map was clearer and biologically more interpretable, with a Rand index of 0.62 compared to 0.15 for the drug sensitivity profiles (Fig. 3A).

**Figure 3.**
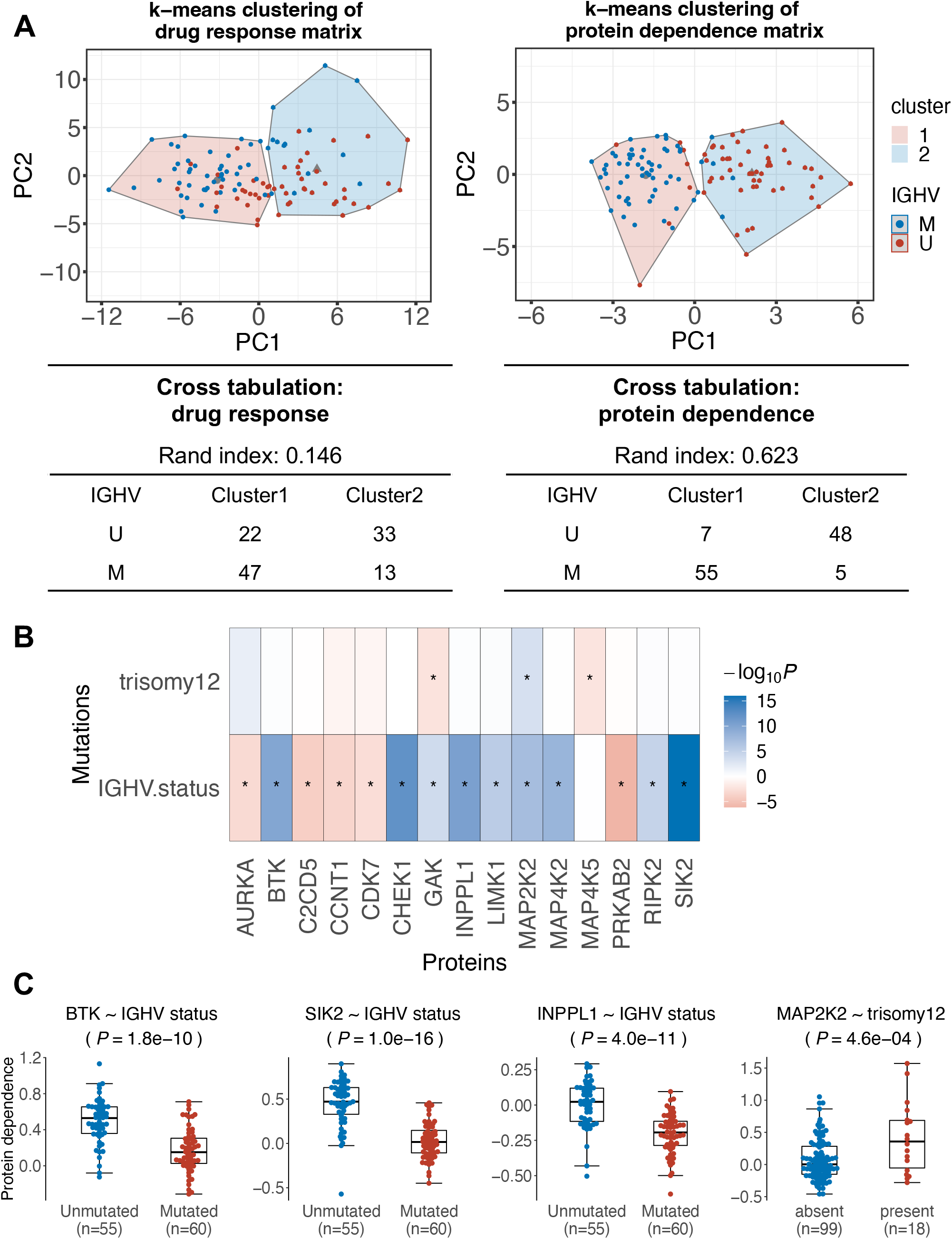
Results of protein dependence inference on primary CLL samples. (**A**) PCA visualization of the CLL samples according to protein dependence matrix (left) and drug sensitivity matrix (right). The points are colored by IGHV mutational status (mutated: blue, unmutated: red), and the results of k-means clustering (k=2) is indicated by the shading. The cross-tabulation and Rand Indices show that the protein dependence matrix-based clustering is more consistent with IGHV mutational status, a known strong stratifying factor in CLL biology (Rand Index = 0.623), than the raw drug sensitivity matrix (Rand Index = 0.146). (**B**) A heatmap showing the -log10(P-values) with signs determined by direction of fold changes of the associations between mutational background of the cell lines and protein dependence coefficients (Student’s t-test). Blue: associations with higher dependence coefficients in trisomy 12 positive / U-CLL; red: higher dependence coefficients in Trisomy12 negative / M-CLL; stars indicate the associated pass 10% FDR control. (**C**) Examples of associations, visualized in beeswarm plots: Association between IGHV mutational status and SIK2 / LYN / BTK dependence and association between trisomy 12 and MAP2K2 dependence. Association testing was performed using Student’s t-test (two-sided, with equal variance) and Benjamini-Hochberg correction was applied.

We looked more specifically into the associations identified between genetic features of CLL patients and the protein dependence coefficients. Most associations were found between IGHV status and protein dependence coefficients (Fig. 3B): as previously described, U-CLL samples showed a higher dependence on BCR signaling components like BTK, LYN, or MAPK/ERK pathway components (Fig. 3 B, C) (Dietrich *et al*, 2017; Guo *et al*, 2016). The inferred protein dependence coefficient of the Salt-inducible kinase 2 (SIK2) also showed a significant association with the IGHV mutational status (Fig. 3 B, C). Notably, several studies suggest SIK2 (and other Salt-inducible kinases) playing a role in the metabolism of B-cell lymphoma (Nagel *et al*, 2010) and AML (Tarumoto *et al*, 2018).

Trisomy 12 - an important cytogenetic driver presents in around 15% of CLL (Döhner *et al*, 2000) - was associated with higher dependence on MAP2K2 (Fig. 3B, C). This association recapitulates a previously reported relationship between trisomy 12 and the MAPK/ERK pathway (Dietrich *et al*, 2017).

#### Protein dependence inference uncovers increased dependence of U-CLL on the cell cycle checkpoint pathway

Having validated the CLL protein dependence map with several associations to known molecular stratifiers of CLL, we looked into novel patterns. The most notable finding was the higher dependence of U-CLL on the checkpoint kinase CHEK1 (Fig. 4A) - a major component of the Chk signaling pathway, compared to M-CLL. A previous study reported higher sensitivity of U-CLL to several CHEK1 inhibitors and hypothesized that this was due to off-target effects of the CHEK1 inhibitors towards components of the BCR-pathway, which are known to have differential dependence between U-CLL and M-CLL (Dietrich *et al*, 2017). DepInfeR enabled us to further disentangle the differential effect of CHEK1 inhibitors and suggests that it is indeed due to higher dependence of U-CLL on the CHEK1 protein, rather than being due to off-target effects of the used CHEK1 inhibitors. Based on the kinobeads dataset, 4 out of 7 CHEK1 inhibitors (kd < 1000nM) also target BCR pathway components (BTK, SYK, LYN) (Fig. 4B). When looking at the effect of the other three drugs, we found a significant association with the IGHV mutational status for two (Rabusertib, MK-8776) of the three CHEK1 inhibitors without any BCR pathway off-targets (Fig. 4C). The third CHEK1 inhibitor (PF-3758309) targets 54 different kinases and binds CHEK1 only very moderately (Kd = 210.19 nM). In combination with our DepInfeR results, this indicates that the differential effect of CHEK1 inhibitors is due to the inhibition of CHEK1 itself and not due to the BCR pathway off-target effect of these drugs.

**Figure 4.**
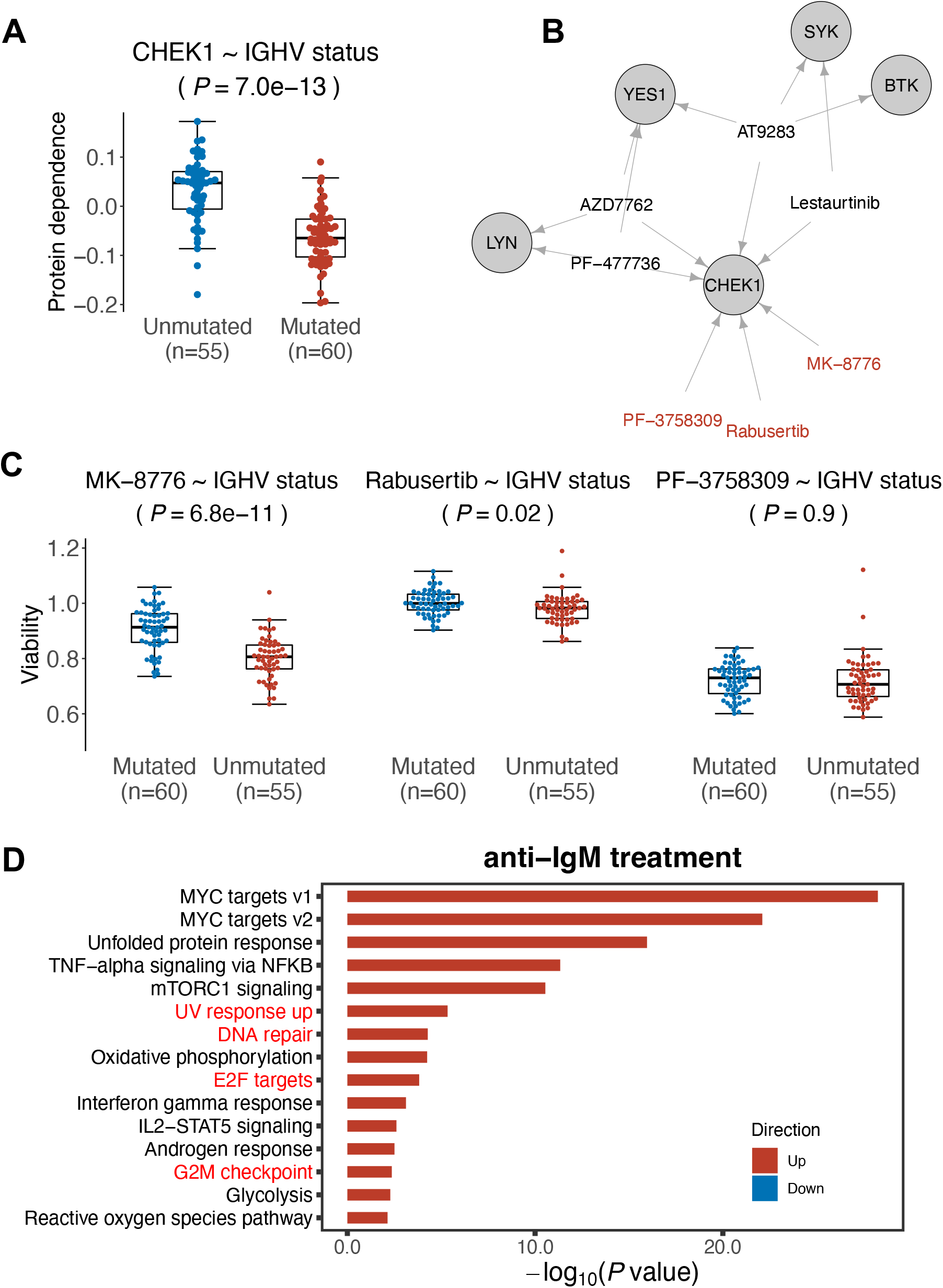
Association of CHEK1 dependence with IGHV mutational status in CLL. (**A**) Beeswarm plot of the CHEK1 protein dependence values in all samples, visualizing the increased dependence on CHEK1 in U-CLL tumors. Association testing was performed using Student’s t-test test (two-sided, with equal variance) and Benjamini-Hochberg correction was applied. **(B)** A network illustration of the off-target effect of CHEK1 inhibitors that involves BCR components (BTK, SYK, YES1 and LYN). Only the high confidence pairs in the Kinobeads dataset are considered. The drugs that only target CHEK1 are colored in red. (**C**) Beeswarm plots showing the effect of three CHEK1-specific inhibitors in U-CLL and M-CLL samples. Association testing was performed using Student’s t-test test (two-sided, with equal variance) and Benjamini-Hochberg correction was applied. (**D**) Differentially expressed hallmark gene sets in CLL cells after BCR pathway stimulation through anti-IgM treatment (ArrayExpress ID: E-GEOD-39411). Highlighted upregulated hallmark gene sets are associated with DNA-damage response and cell cycle checkpoint.

The association of U-CLL to CHEK1 pathway was further corroborated by an analysis of differential gene expression between M-CLL and U-CLL patient samples from the data of *Ferreira et al*. (European Genome-phenome Archive; http://www.ebi.ac.uk/ega/, EGAS00001000374). Amongst the most upregulated pathways in U-CLL (Supplementary Figure S8) were DNA-damage response- and cell cycle checkpoint-related pathways (G2/M checkpoints, DNA repair, E2F transcription factor targets, UV response) (Verlinden *et al*, 2007), and we found several genes upregulated that are directly involved in CHEK1-mediated checkpoint control (e.g. *CDC25A, CDK1, CCNE1*) (Supplementary Figure S9).

Moreover, we found that BCR stimulation by anti-IgM antibodies in CLL cells led to a significant upregulation of pathways related to DNA-damage response and cell cycle checkpoint, including UV response, DNA repair, E2F transcription factor targets, and G2/M checkpoints (Fig. 4C). This finding is in line with previous studies showing that CHEK1 plays a role in the survival of BCR-activated and developing B-cells (Schoeler *et al*, 2019; Schuler *et al*, 2017).

Taken together, our results suggest a stronger dependence of U-CLL on CHEK1-mediated checkpoint control. This might be either a direct effect of the increased BCR pathway activity in U-CLL or a result of increased replication stress in this more aggressive CLL subtype. Targeted CHEK1 inhibition may therefore be especially beneficial for U-CLL.

## Discussion

We developed a computational framework, DepInfeR, that integrates drug response assays and drug-protein affinity data to infer the sample-specific protein dependencies, which reflect how much the survival of the cancer cells (from a specific cell line or patient sample) depends on a certain protein.

We contrast some of the methodological choices of DepInfeR to a recent proposal by (Wang *et al*, 2020), who pursued a similar aim by taking a drug-protein binding graph using the mean effect of all drugs connected to a given protein in the graph as a measure of dependence on it (termed *target addiction score* in their study). In contrast, our approach takes a more quantitative approach by representing protein-drug affinities with a matrix whose elements take values between 0 and 1, and if quantitative measurements such as *K*_*d*_ values are available, uses these by mapping them to the range [0,1] with a sigmoid function. Moreover, we use a set of linear equations to relate protein dependencies (not directly observed), drug-protein affinities (data) and cell viabilities (data) and use multivariate regression with regularization to solve for the quantities of interest. The regularization step provides sparsity—many of the estimated effects are zero—, which can facilitate interpretation and reduce noise, and is consistent with the notion that not every drug-protein binding event will contribute to the observed phenotype of the drug. Our approach stops short of more realistic biophysical modelling of the drug-protein binding kinetics: this would be an interesting direction for future research.

We validated our approach by showing that it could accurately recapitulate known protein dependencies of different cancer entities as well as of their molecular subtypes, both in data from cancer cell line compendia and from primary patient tumor samples. We then discovered a novel finding, the high dependence of CLL with unmutated IGHV status (U-CLL) on the Checkpoint kinase CHEK1. This protein is an important component of the cellular DNA-damage and cell cycle checkpoint response and has been targeted in cancer therapeutics, especially for MYC-expressing cancers (Höglund *et al*, 2011; Ferrao *et al*, 2012). The critical role of CHEK1 in rapidly proliferating, developing, and BCR-activated normal B-cells was previously noted (Schoeler *et al*, 2019; Schuler *et al*, 2017). The drug screen by *Dietrich et al. 2017* found high sensitivity of U-CLL to CHEK1 inhibitors, however, due to the promiscuity of CHEK1 inhibitors, including towards other kinases known to be important in CLL, the interpretations remained ambiguous. However, our results, in combination with these previous observations, suggest a model where U-CLL tumor cells, which often exhibit increased tonic or antigen triggered BCR signaling, experience more stress-related signals due to higher metabolic activity or increased replication stress. This might prime cancer cells towards a highly activated CHEK1-mediated cell cycle checkpoint control (which remains active in *in vitro* culture). As a result, U-CLL cells are more vulnerable to CHEK1 inhibition, driving these cells towards apoptosis. Our results suggest targeted CHEK1 inhibition as a potential therapeutic strategy in the U-CLL patients.

The inference accuracy of our method relies on the size and quality of both the drug-protein profiling matrix and of the ex-vivo tumor-drug response matrix. Generally, any increase in the number of drugs, tumor samples or proteins is beneficial. The data preprocessing steps, such as the transformation of drug-protein binding affinity matrix and the summarization of the drug dose-response values, may also impact the final inference results. Therefore, it requires further research to answer important statistical questions of identifiability and uncertainty in the estimated protein dependence map.

While our present analyses were limited to kinases and kinase inhibitors, we expect that the approach would be analogously applicable should protein-drug binding matrices for other protein classes become available.

In conclusion, we present a method to infer protein dependence maps from drug sensitivity data. Among the possible uses of this method, we envisage both, the discovery of novel disease stratifications by their characteristic dependencies on specific proteins, and improved understanding of existing disease stratifications in terms of differential protein dependencies. Nevertheless, such stratifications gain potential therapeutic value by being directly associated with druggable targets.

## Acknowledgments

J.Lu. and W.H. were supported by the German Federal Ministry of Education and Research (CompLS Project MOFA under grant agreement 031L0171A, and SMART-CARE under grant agreement 031L0212E). T. Zenz was supported by the CRPP “Next Generation Drug Response Profiling for Personalized Cancer Care “, the Swiss Cancer Research foundation (KFS-4439-02-2018), and the Monique-Dornonville-de-la-Cour Stiftung. J.K. was supported by postdoctoral fellowships from the Sigrid Juselius Foundation and the Finnish Cultural Foundation.

## Author contributions

Conceptualization W.H., J.Lu; Experimental design: T.Z., J.K.; Software, formal analysis, visualization: A.B., J.Lu; Methodology and experimental investigation: J.Lewis, K.P.; Writing the manuscript: A.B., J.Lu, W.H., T.Z., J.K.; Data curation: J.Lu, K.P.; Resources: J.Lu, A.B.

