## Supplementary Materials for "Inferring tumor-specific cancer dependencies through integrating ex-vivo drug response assays and drug-protein profiling"

### 1 Supplementary Figures

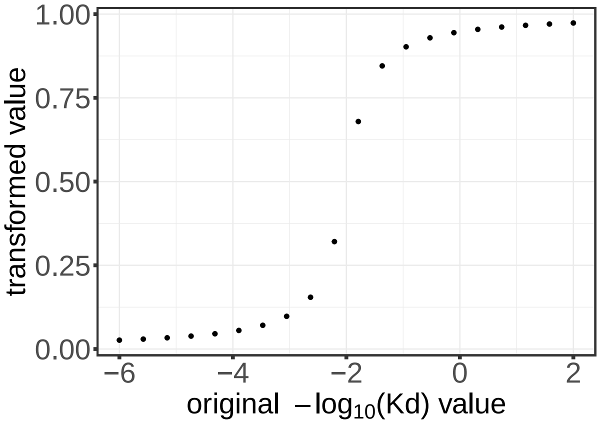


Figure S1: Plot of the function, $f(x) = \frac{arctan((x+2)*3)+\frac{\pi}{2}}{\pi}$, that is used to transform the $-\log_{10}(Kd)$ values from Kinobeads assays.

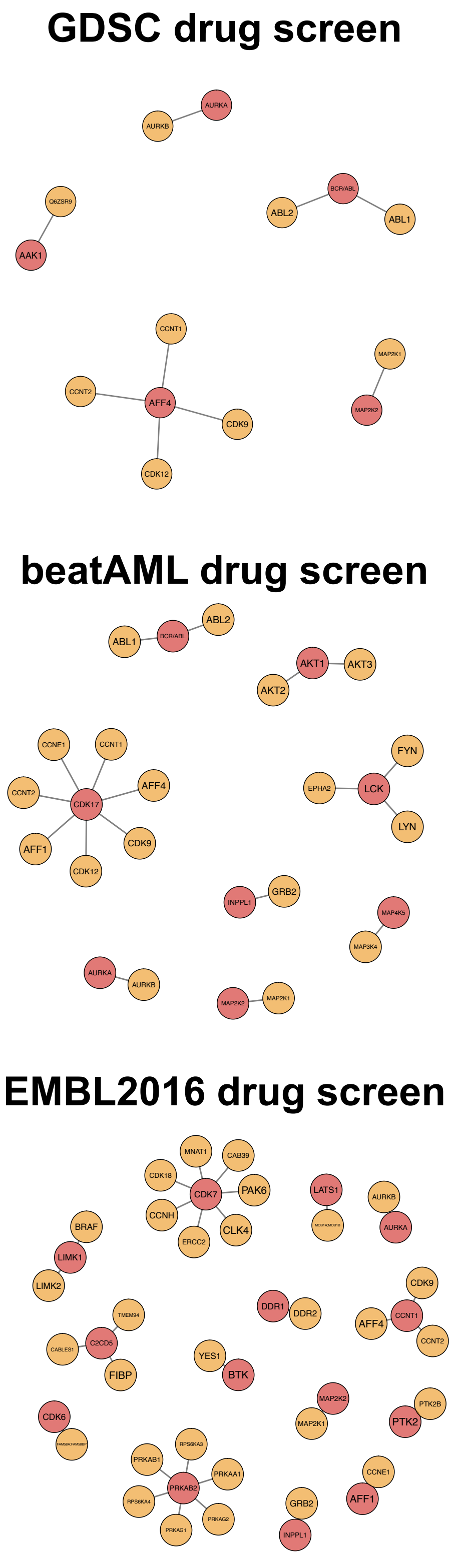


Figure S2: Network plots showing the clusters of kinases that have similar drug binding profiles (cosine similarity $> 0.8$). For each drug screen dataset, only the kinases that were selected by our protein dependence inference algorithm and share similar drug binding profiles with other kinases are shown here.

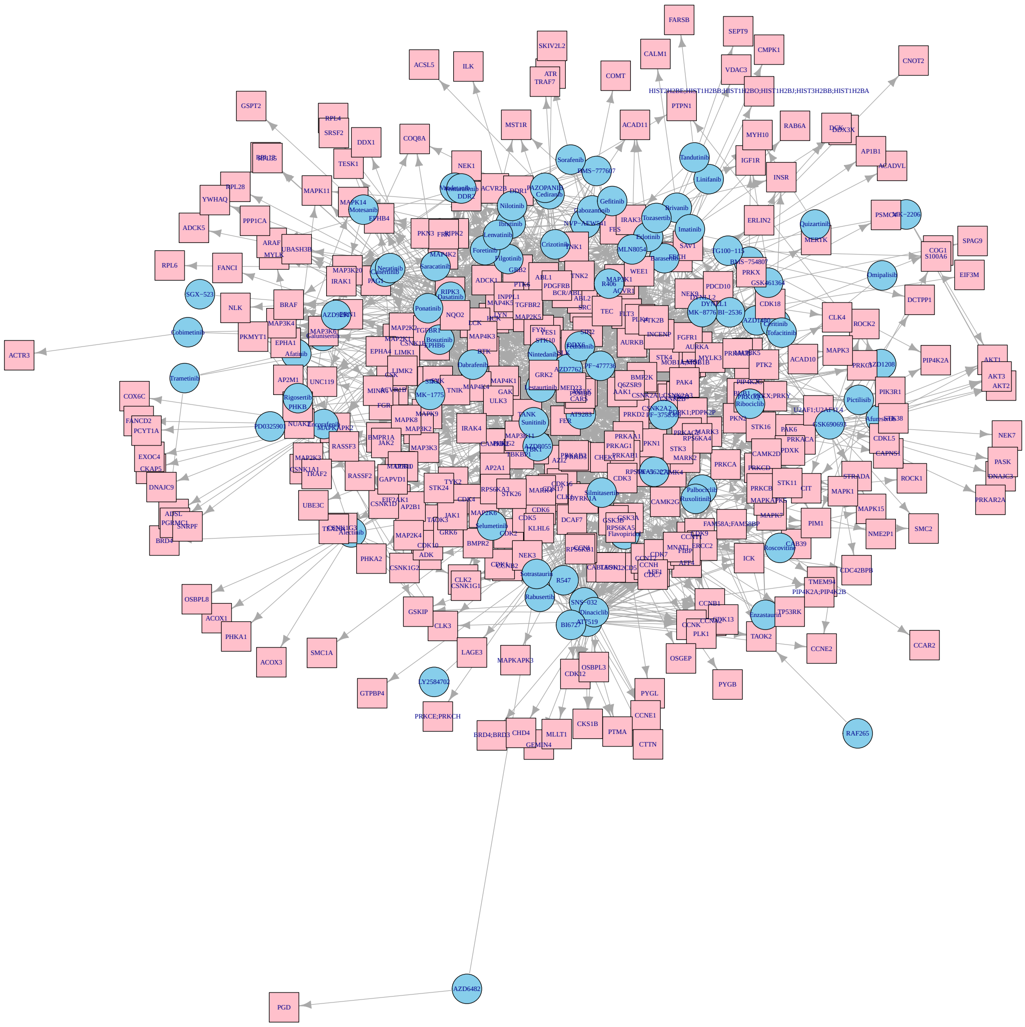

Figure S3: A network plot showing the drug-kinase binding landscape measured by the Kinobeads assay. Only the high confidence drug-kinase pairs are included. Drugs are shown as blue circles and kinases are shown as red squares. No clear structure can be observed directly from the drug-kinase network.

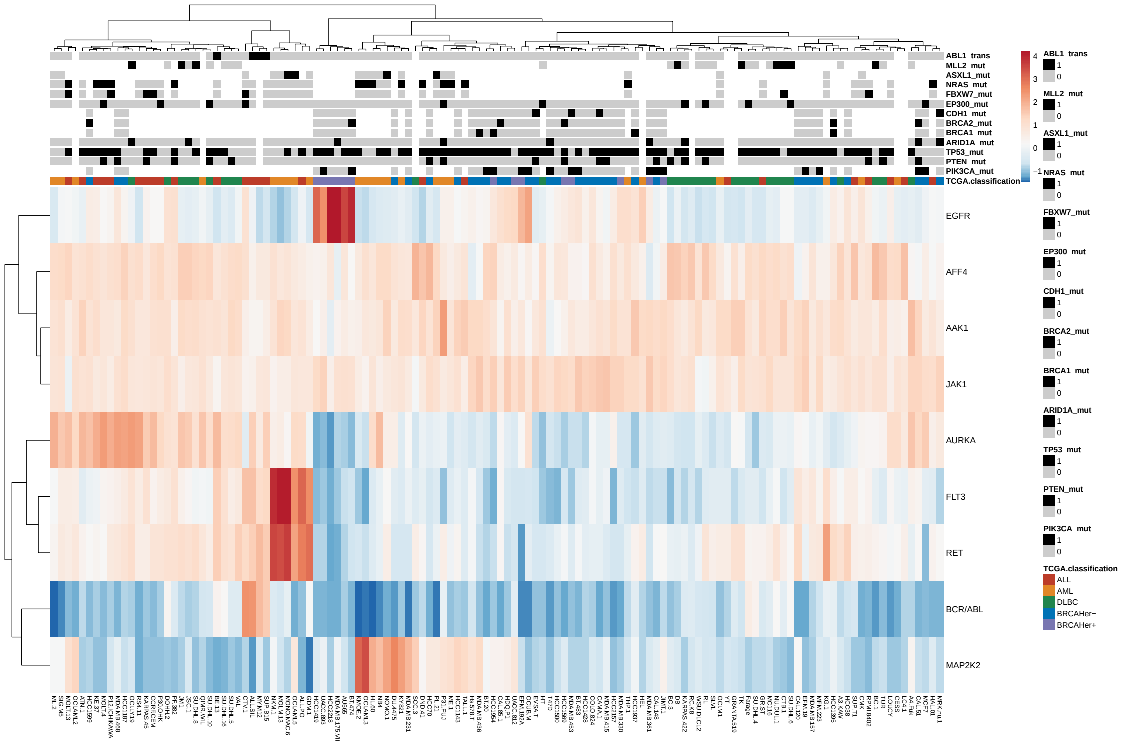


Figure S4: Heatmap visualization of the protein dependence values for selected kinases, inferred from the GDSC drug screen dataset. Red indicates high dependence value and blue indicates low dependence value. In the column annotations, cell lines with certain genomic variations are colored by black. Columns are ordered by hierarchical clustering based on euclidean distance.

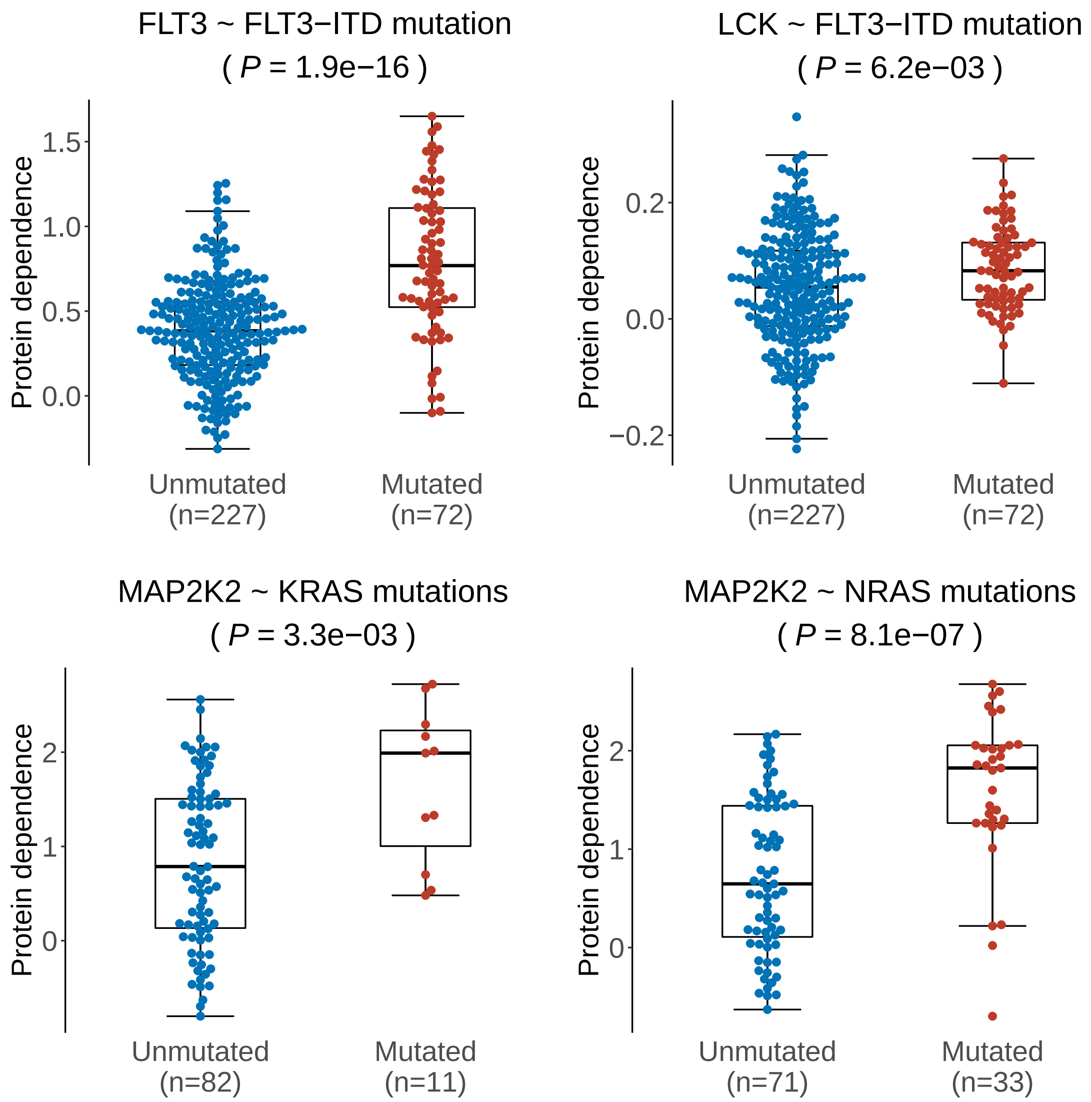


Figure S5: Boxplots showing the significant associations between the genomic variations and the protein dependence values inferred from the beatAML drug screen dataset.

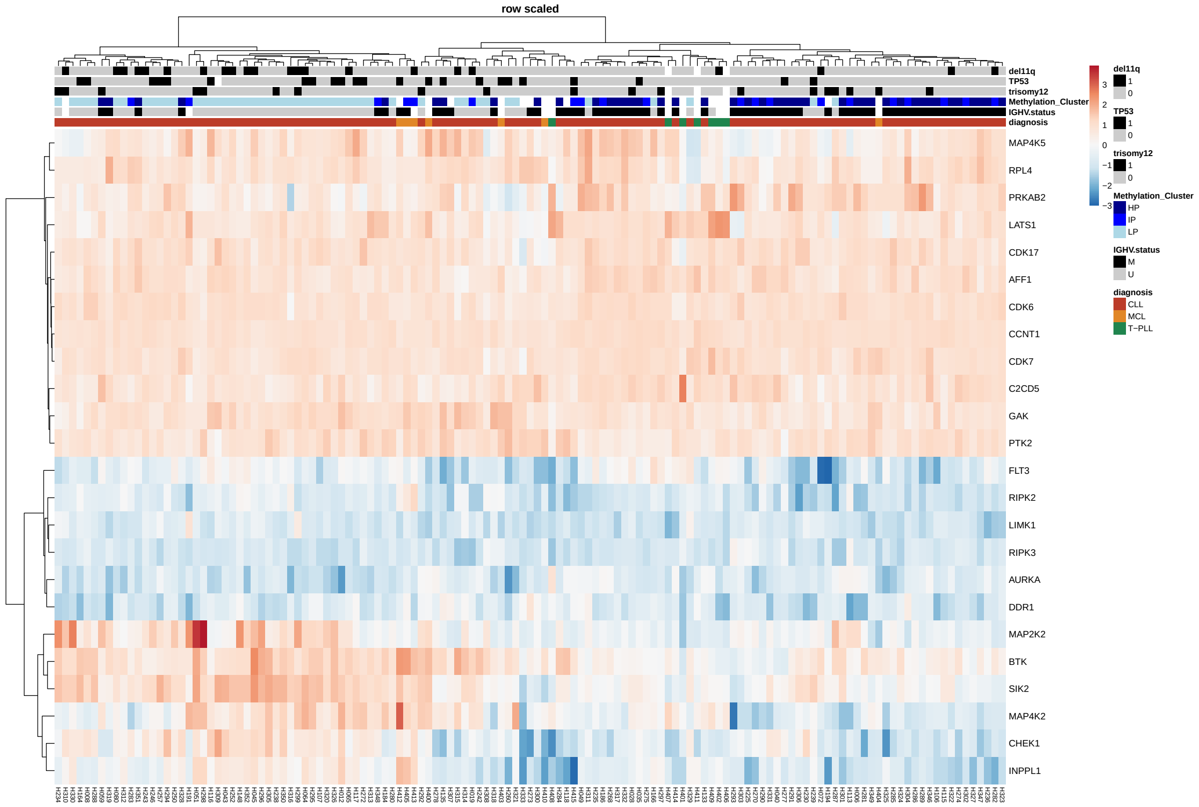


Figure S6: Heatmap visualization of the protein dependence values for selected the kinases inferred from the EMBL2016 drug screen dataset. Red indicates high dependence value and blue indicates low dependence values. The column annotations of the heatmap indicate the disease types, genomic variations and epigenetic subtypes of the primary blood cancer samples. Columns are ordered by hierarchical clustering based on euclidean distance.

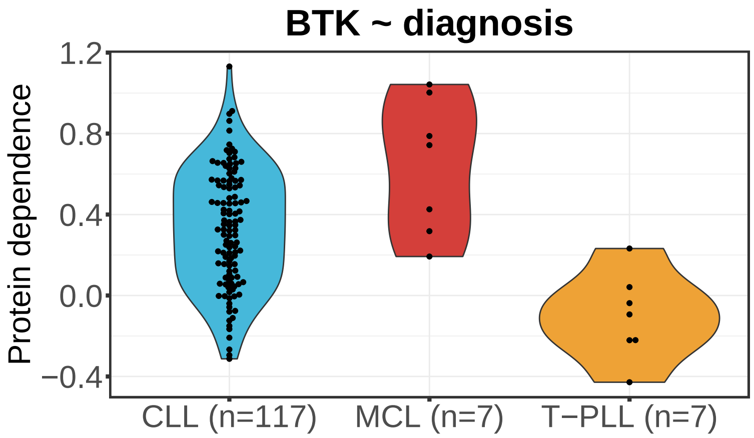


Figure S7: Association between the inferred dependence on BTK and the disease types, based on the EMBL2016 drug screen dataset.

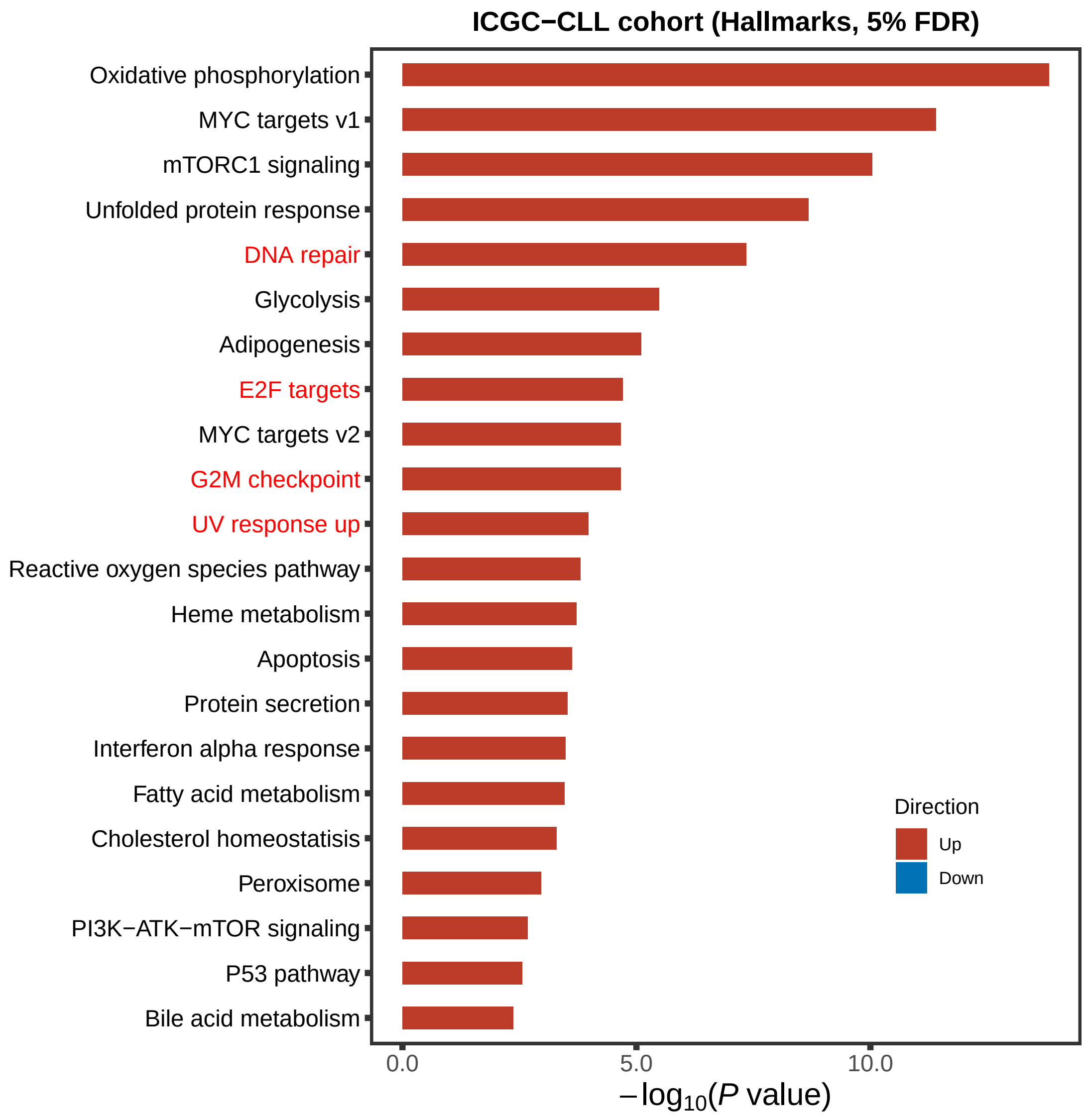


Figure S8: Enrichment analysis for genes differentially expressed between IGHV mutated and unmutated CLL samples in the ICGC-CLL cohort (EGA accession ID: EGAS00001000374). ”Up” (colored by red”) indicates the gene sets enriched for the genes up-regulated in U-CLL samples compared to M-CLL samples. The enrichment analysis was perform by using CAMERA against the Cancer Hallmark gene sets from the Molecular Signature Database (MSigDB). Only the gene sets passed 5% FDR are shown. The gene sets that have been shown to be related to CHEK1 functions are colored in red.

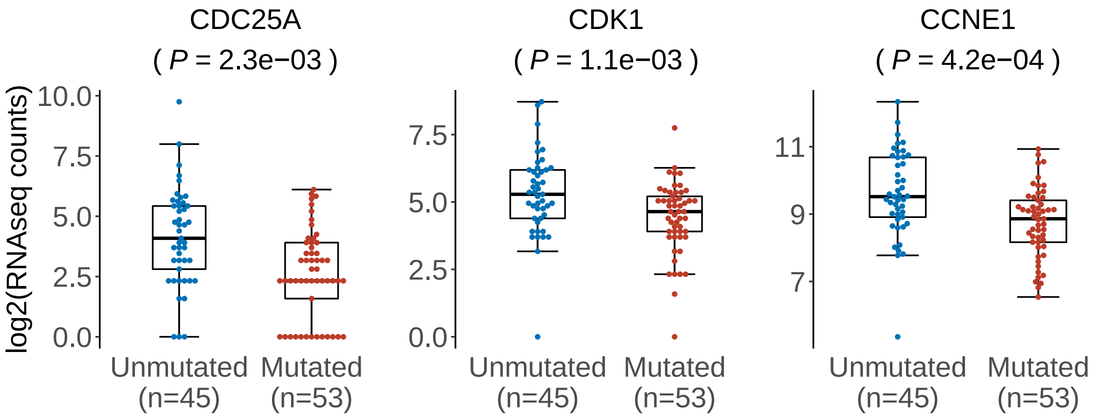


Figure S9: Differential expression of three downstream genes regulated by CHEK1 pathway in IGHV unmutated and mutated CLL samples from the ICGC-CLL cohort. $P$ values were calculated by DESeq2.
